# CaSee: A lightning transfer-learning model directly used to discriminate cancer/normal cells from scRNA-seq

**DOI:** 10.1101/2022.02.10.480003

**Authors:** Yuan Sh, Caixia Guo, Fanghao Shi, Fei Jia, Zhiyuan Hu, Xiuli Zhang

## Abstract

Single-cell RNA sequencing (scRNA-seq) is one of the most efficient technologies for human tumor research. However, data analysis is still faced with some technical challenges, especially the difficulty in efficiently and accurately discriminate cancer/normal cells in the scRNA-seq expression matrix. In this study, we developed a cancer/normal cell discrimination pipeline called pan-cancer seeker (CaSee) devoted to scRNA-seq expression matrix, which is based on the traditional high-quality pan-cancer bulk sequencing data using transfer learning. It is compatible with mainstream sequencings technology platforms, 10x Genomics Chromium, Smart-seq2, and Microwell-seq. Here, CaSee pipeline exhibited excellent performance in the multicenter data evaluation of 11 retrospective cohorts and one independent dataset, with an average discrimination accuracy of 96.69%. In general, the development of a deep-learning based, pan-cancer cell discrimination model, CaSee, to distinguish cancer cells from normal cells will be compelling to researchers working in the genomics, cancer, and single-cell fields.

## Introduction

As scRNA-seq technologies have been rapidly developed in recent years, we could easily study the characterization, development stage and specific stage of cell heterogeneity in ultra-high throughput sequencing data^1,2^, especially in the research on the occurrence and development of cancer^3,4^. However, large-scale data analysis is still faced with a huge and pivotal challenge of accurately discriminating cancer/normal cells in the scRNA-seq expression matrix. At the single-cell level, only by accurately discriminating cancer/normal cells in the matrix can we further analyze intratumoral and intertumoral heterogeneity^5^.

There are two main traditional methods for the discrimination between cancer and normal cells: 1) cell clustering methods based on marker gene^6^. 2) CNV detection methods^7^. In the main cancer cell discrimination method based on marker genes, cell types are manually annotated based on database or software, such as CellMarker^8^, CancerSEA^9^ and SingleR^10^ (https://github.com/dviraran/SingleR). However, it is difficult to use these databases or automatic annotation software to discriminate cancer/normal cells in the scRNA-seq expression matrix.

In contrast, it is relatively easy to use CNV-based cell discrimination method. CNV detection methods can be applied under a vital hypothesis that most cancer cells have CNVs (> 85%)^11^. However, research has shown that somatic copy number alterations (SCNAs) are widely present in normal human tissues ^12,13^. More significantly, normal cells have a higher frequency of CNVs in the tumor microenvironment (TME) than in the normal environment^13^. Therefore, only applying CNV detection methods cannot accurately discriminate between cancer and normal cells. In addition, the two commonly used CNV detection methods, namely, inferCNV^4^ and CopyKAT^14^, based on statistical methods, have limitations, that is, they cannot use graphics processor (GPU) for accelerated calculation and can only be processed in central processing unit (CPU). Therefore, the application of both inferCNV and CopyKAT is time-consuming, especially in a large data set, and they function at the population level. For example, inferCNV uses stromal or immune cells as references to observe candidate cancer cells and then selects a suitable threshold of the output value according to the distribution of the whole dataset to identify the “identity” of each cells^7^. Besides, CopyKAT infers aneuploidy according to the statistical population, so it cannot directly respond to the data with high purity of cancer cells or shows no obvious change in the CNV level^14^. In closing, during the application of inferCNV and CopyKAT, the accuracy of results will change with background data, so both tools cannot be directly used to discriminate cancer/normal cells in the scRNA-seq expression matrix.

With the rapid development of deep learning technology, it is common to use deep learning methods, such as scDeepSort^15^, a cell annotation tool based on weighted graph neural network (GNN), and DISC^16^, a semi supervised deep learning model that can infer the gene structure and expression masked by drop-out, to explore scRNA-seq expression matrix. These emerging deep learning analysis methods provide us with a novel idea in scRNA-seq data mining. At the algorithm technology level, the strength of deep learning lies not only in its ability to achieve accurate sample classification based on big data, but also in its excellent “transfer learning” ability^17–20^ to learn in a specific data space, and then apply the “knowledge” in subspace or similar data space^17,18^. At the hardware equipment level, the increasingly mature GUP hardware technology greatly shortens the time of deep learning training and prediction and greatly improves the efficiency of analysis. At the dataset level, with the rapid development of NGS omics technology, the number of high-quality bulk data collected in the past was extremely large such as TCGA, which provides a large number of mining conditions for studying scRNA-seq data. At the sequencing technology level, scRNA-seq data are homologous with bulk sequencing data, and many single-cell level research conclusions can be reproduced in bulk data^21–23^. In this research, we integrated the bulk data and established a pan-cancer cell discrimination model (Pan-cancer seeker, CaSee) to discriminate cancer/normal cells in bulk data in the scRNA-seq expression matrix. Subsequently, we evaluated the discrimination performance of the model using a variety of methods and proved that CaSee could discriminate cancer/normal cells in scRNA-seq expression matrix, quickly and accurately.

## Results

### Overview of CaSee pipeline

CaSee requires scRNA-seq raw count matrix or the count matrix which only contains candidate cancer cells. In CaSee pipeline, data were obtained from TCGA and GTEx as “candidate reference data”, which involved 18 cancer types, containing 7,114 tumor samples and 6,774 healthy samples (n=13,888 in total; Fig.1A top; Supplementary Table 1). Users could directly input raw scRNA-seq count matrix for subsequent discrimination procedure. The identification module, named “cell exclusion”, for candidate cancer cells has been embedding in the CaSee (see configs-file). After CaSee automatically identified all the candidate cancer cells, pipeline start. (Fig.1B top). To generate the similarity sample space, candidate cancer matrix intersected with candidate reference data to output “reference data” (Fig.1A, b bottom).

Then the “reference data” were split into three parts, training dataset, validation dataset and test dataset. Next, the training dataset was input into the training loop and the validation dataset to select the best model parameter. However, CaSee is a kind of capsule neural network^24–26^ belonging to the 2D convolutional neural network, which is widely used in image classification tasks^24,26,27^. In addition, since RNA-seq data are 1D tensors that are not supported by 2D convolutional neural networks. Therefore, we designed a tensor conversion layer as the input layer of the model. The vector in each sample (each column of RNA-seq expression matrix) passing the layer was converted into a feature map (default 28 × 28), and each pixel of the feature map was an abstract gene module composed of multiple genes (Fig.1C FC). Next, the following network structure expand feature map into a capsule (Fig.1C Mapping Encoder). Then the capsule would pass through the capsule encoder, followed by dynamic routing^28^ (Fig.1C Capsule Encoder & Dynamic Routing), and due to the unique nature of the capsule neural network, the relative position relationship between each abstract gene module was preserved (Fig.1C Capsule)^28^. At the end of the model, we set up the decoding capsule module to retain the original information of the data as much as possible while ensuring the classification accuracy, so as to ensure the complete cancer-normal space and the success of transferring model^26^. Finally, the optimal model qualified in the test dataset (accuracy>0.95) was transferred into the candidate cancer cells expression matrix to discriminate cancer/normal cells (Fig. 1C).

**Fig.1.**
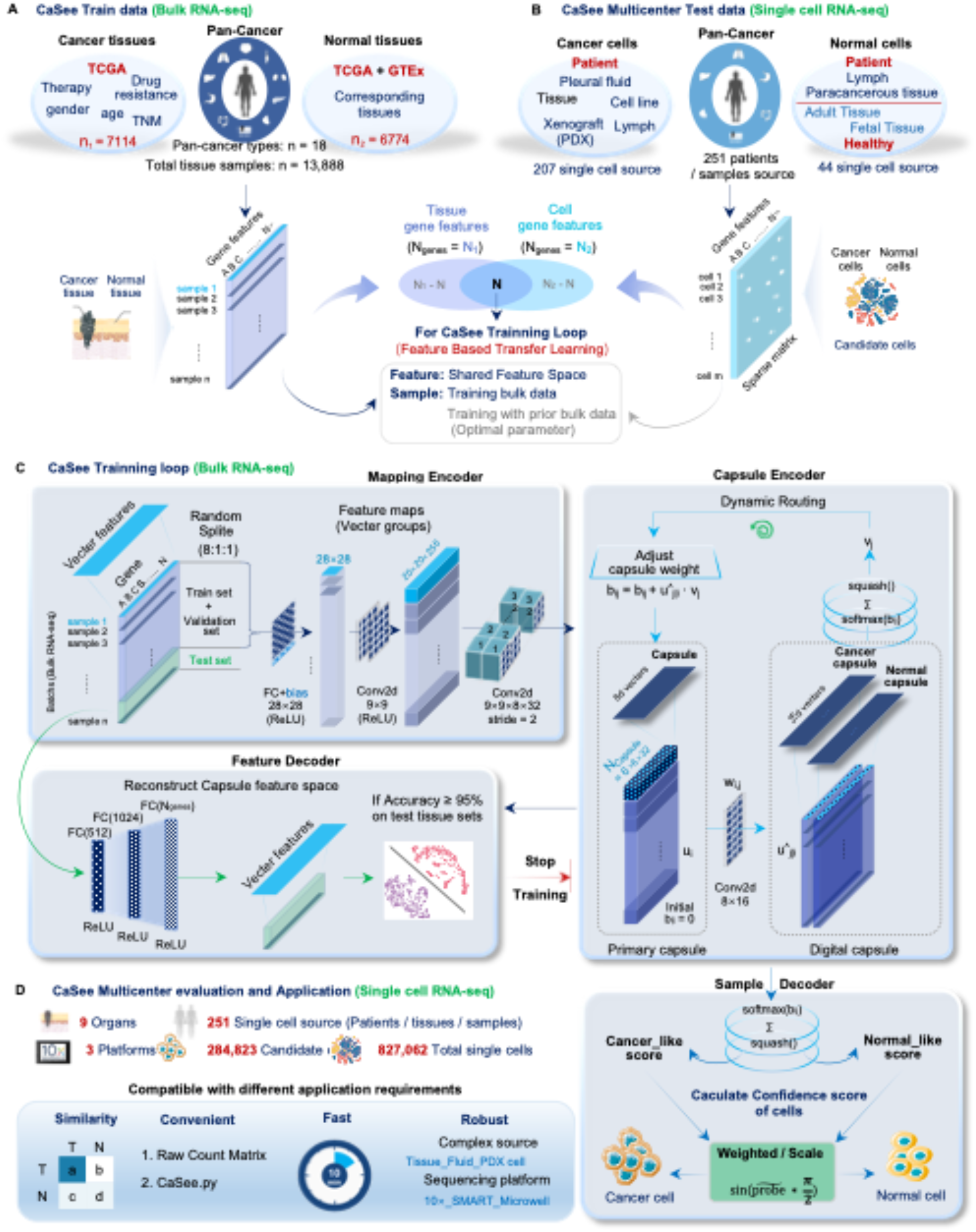
Introduction and overview of the CaSee Model workflow. A, (top) Summary of “candidate reference data” which is composed of TCGA and GTEx (detail information see supplementary table1). B, (top) 251 samples were used to evaluate the performance of CaSee Model, including 207 Cancer source and 44 Normal sources (detail information see supplementary table2). A, B (bottom) Gene features were overlapped between candidate reference data and single-cell RNAseq, and the output bulk data was considered as “reference data” which has a particular dataspace similar to target scRNA-seq count matrix. C, (top) CaSee model training loop. First, the reference data was shuffled and randomly split into training: validation: test (8:1:1); second, go to the feature map extraction layer convert 1D tensor to 2D tensor; third, each feature map in turn through Capsule layer, Dynamic layer, and Decoder layer, finally output decoder matrix and predict. (left bottom) Only when the final output model can accurately predict testing data, (right bottom) can the model transfer to scRNA-seq data and there is tow out predict result (scale probability and weighted probability). D, Summary of the model robust evaluation and the advantages. FC, full connected layers.

Considering the sparsity of scRNA-seq space^29^ and sample heterogeneity, CaSee provides two strategies to adjust the output probability. The first one is scale_predict that directly compares normalized absolute probability of cancer or normal cells (see Methods). The second one is weight_scale_predict that is weighted scale_predict (see Method). Additionally, CaSee also functions in a multi-cross mode, under which users can train more than one model to delineate input data. It will integrate all model’s output files and use the cumulative and binomial distribution test to get each cell’s p-value. Benjamini-Hochberg (BH) adjustment method was adopted to control false discovery rate (FDR) and select adjusted *p-values*<0.05 as the threshold to discriminate cells (see Method). We used 11 retrospective cohorts^6,30– 37^ and one independent dataset to evaluate the robustness of CaSee. All data contained 9 tissue types, 251 data sources and 3 platforms, and there were a total of 827,062 cells including 284,823 candidate cells (Fig. 1D, Supplementary Table 2).

### Evaluation of CaSee performance

Firstly, the performance of CaSee evaluated in high-purity scRNA-seq data (n = 16), including SKCM xenograft cell, cell line, healthy adult samples, and fetal samples, average accuracy is 99.81% (Fig. 2A Supplementary Table 2, 3). Then we selected four representative samples (cancer source = 2, normal source = 2) and statistics of cancer probability (Normal probability=1-Cancer probability) distribution in each data. In the cancer source data, the probability of cancer sources closes to 1 and the counterpart closes to 0. The results preliminarily demonstrated that the CaSee model had high accuracy in discriminating cancer/normal cells (Fig. 2B).

**Fig. 2.**
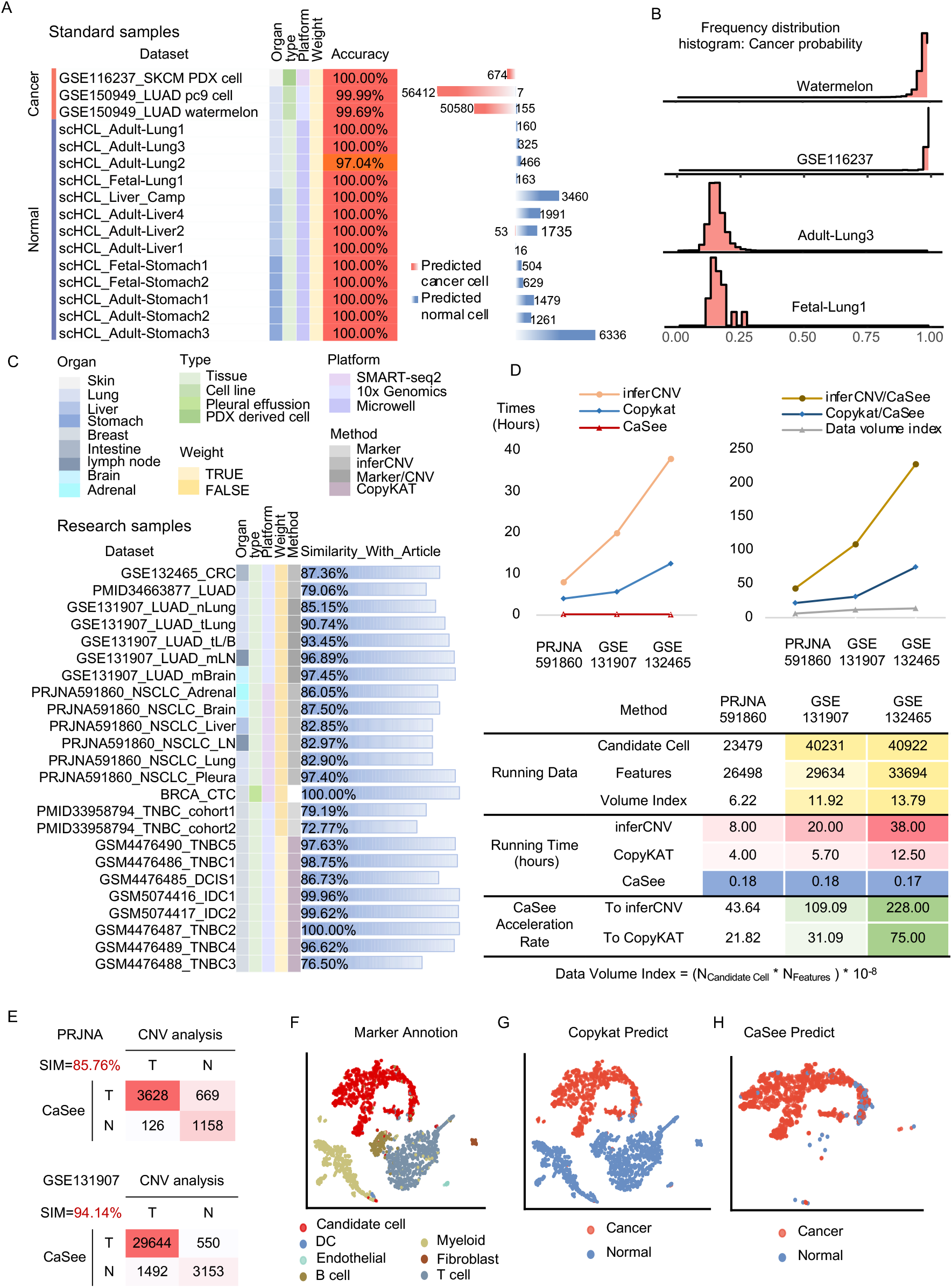
Comparison of CaSee and copy number variation method. A, High purity cell source datasets consider as the standard samples used to evaluate the accuracy and performance of CaSee, which is contained SKCM xenograft cell, cell line, healthy adult sample, and fetal sample. B, Frequency distribution histogram of the probability of cancer cells output by CaSee, Watermelon and GSE116237 are cancer origin, as well as Adult-lung3 and Fetal-1 are normal origin, (Normal probability=1– Cancer probability). C, Multicenter data evaluation in 11 of independent datasets and one Circulating tumor cell (CTC) datasets. D, (top right) The running time of the three methods in different datasets; (top left) Volume index and running time ratio (inferCNV/CaSee and CopyKAT/CaSee) in three datasets; (bottom) Heatmap summary of running time performance, exact value in Supplementary Table 4. E, Confusion matrix and similarity of article label. F-H, Similarity between CopyKAT and CaSee, tSNE plot in different method result. F, using markers to define cell cluster, (all markers in Supplementary Fig. 1 bottom Dotplot). G, CopyKAT predict result to define cell cluster. H, CaSee predict candidate cells.

Considering the actual application scenarios of the model, we evaluated the consistency of discrimination results using CaSee with those using traditional methods in the candidate cancer cells in different research (Supplementary Table 2). The results revealed that in many studies, the results of CaSee were highly consistent with those defined by researcher based on traditional methods (Fig. 2C). In addition, we also compared the running time of inferCNV, CopyKAT and CaSee. Obviously, with the growth of the volume index of datasets, the running time of inferCNV and CopyKAT surged, while CaSee was almost stable (Fig. 2D). The reason may be that the pipeline construction process of CaSee is only related to the “reference data”, independent of the volume index of scRNA-seq data. The running time of CaSee was significantly faster than inferCNV and CopyKAT, and the average speed up by about 127 times and 43 times, respectively (Fig. 1D bottom, Supplementary Table 4).

Interestingly, the comparison of discrimination results between inferCNV and CaSee showed a completely opposite tendency. Compared with CaSee, the inferCNV method was more inclined to catching normal cells in PRJNA591860^38^ but catching cancer cells in GSE131907^7^ (Fig. 2D). This trend showed statistically significant differences at different biopsy sites (Supplementary Table5). This conclusion indicated that there is no obviously statistical prejudice in the discrimination process of different data by CaSee. To figure out the cause of this phenomenon, we evaluated the performance of CaSee by strictly abiding by the rules set by the researchers. In both PRJNA591860 and GSE131907, inferCNV method was adopted to discriminate cancer/normal in candidate cell clusters, but their identification strategies were different. As mentioned above, inferCNV method cannot directly get the exact “identity” of cells. Therefore, researchers need to manually set a relevant threshold and strategy to classify each “identity” of cells. In the study of PRJNA591860, researchers considered those above the threshold as cancer cells and others as normal cells, while in the GSE131907, research adopted different discrimination strategies. They tagged the cell “identity” according to different biopsy locations and patient’s information combined with the results of inferCNV based on the following rules: if cells with inferCNV result above the threshold from the lesions or the metastasis site will be considered as cancer cells, otherwise, tagged NA; cells with inferCNV result below the threshold from normal tissues will be considered as normal cells, else tagged NA. Therefore, in the research of GSE131907, there are parts of cells did not support the conclusion of the study. Next, we evaluated the similarity between CopyKAT and CaSee, and the results showed that CaSee could maintain a high degree of consistency with CopyKAT (Fig. 2F-G, Supplementary Fig1). In conclusion, CaSee can maintain a high degree of similarity with the two traditional CNV methods, and it provides faster and easier solution.

### Application evaluation of CaSee in exploring lung cancer cell-specific pattern compared with inferCNV

As mentioned above, the inferCNV method produces different results based on different discrimination strategies. This section will try to evaluate the results of different strategies *vs*. CaSee model results from different aspects, with GSE131907 as an example. Here, we obey the rules formulated by the researchers (Article group). However, considering that not all studies can obtain control samples, we retained all the results of inferCNV and considered the undetermined as normal (inferCNV group). What’s more, we define the absolute group, which is the result of CaSee (CaSee group) and completely consistent with that in article (Absolute group). Totally, there were four groups in following evaluation process: 1, Absolute; 2, Article; 3, CaSee; 4, inferCNV.

As noted by Zhang L *et al*., we found the same phenomenon with Maynard Ashley *et al*. Compared to normal cells, cancer cells significantly expressed a greater variety of genes (Fig. 3A)^22,39^. The more genes, the more differential expression genes (DEGs). We then identified DEGs in four groups (cancer *vs*. normal), respectively (cancer *vs*. normal). The statistical results of DEGs (abs (logFC) > 2 and adjusted *p-value* < 0.05) showed that the distribution of DEGs in Absolute group was similar to that in Article group and CaSee group, and the number was larger than that in inferCNV group (Fig. 3B), and the total number of DEGs was 4,451 (Fig. 3C right). Next, we evaluated the consistency of DEGs between each group (see Method). We took Absolute group as the standard. The consistency of DEGs between Article group and CaSee group with the standard was 80.7% and 76.32% respectively, while the consistency of inferCNV was only 46.14% (Fig. 3C left). In addition, we found that the cell annotation distribution in the CaSee group was highly consistent with that in the Article group (Fig. 3D). The results preliminarily showed that the results of the application of only inferCNV could produce some errors in complex data (the errors in GSE131907 are mainly concentrated in PE). Reasonable formulation of discrimination strategy is of great help to the research, but it has strict requirements for data collection. Comprehensively, the CaSee is slightly better than the inferCNV. The result of IPA analysis gave the same supports. In the Absolute, Article and CaSee groups, the activation/inhibition degree of both proliferation-related and apoptosis-related functions was more significant than that in inferCNV group, further indicating that apparent pathway/function differences between cancer cells and normal cells tagged by Article and CaSee groups. This difference is strongly related to cancer or tumor biological signals (Fig.3E). It is worth noting that the inferCNV method is not sensitive to free epithelial cells from pleural effusion, which is an essential reason for a large number of “normal cells” in the inferCNV group, resulting in insufficient cell purity in the inferCNV group. Therefore, compared with the direct use of inferCNV, the discrimination strategy designed by Kim Nayoung et al., was reasonable. Unfortunately, this strategy is difficult to popularize for the following reasons: 1. Not all studies can obtain normal control samples; 2. Different research data need to adopt different thresholds, and appropriate threshold setting is the key. CaSee can provide users with faster, better, and easier solution.

**Fig. 3.**
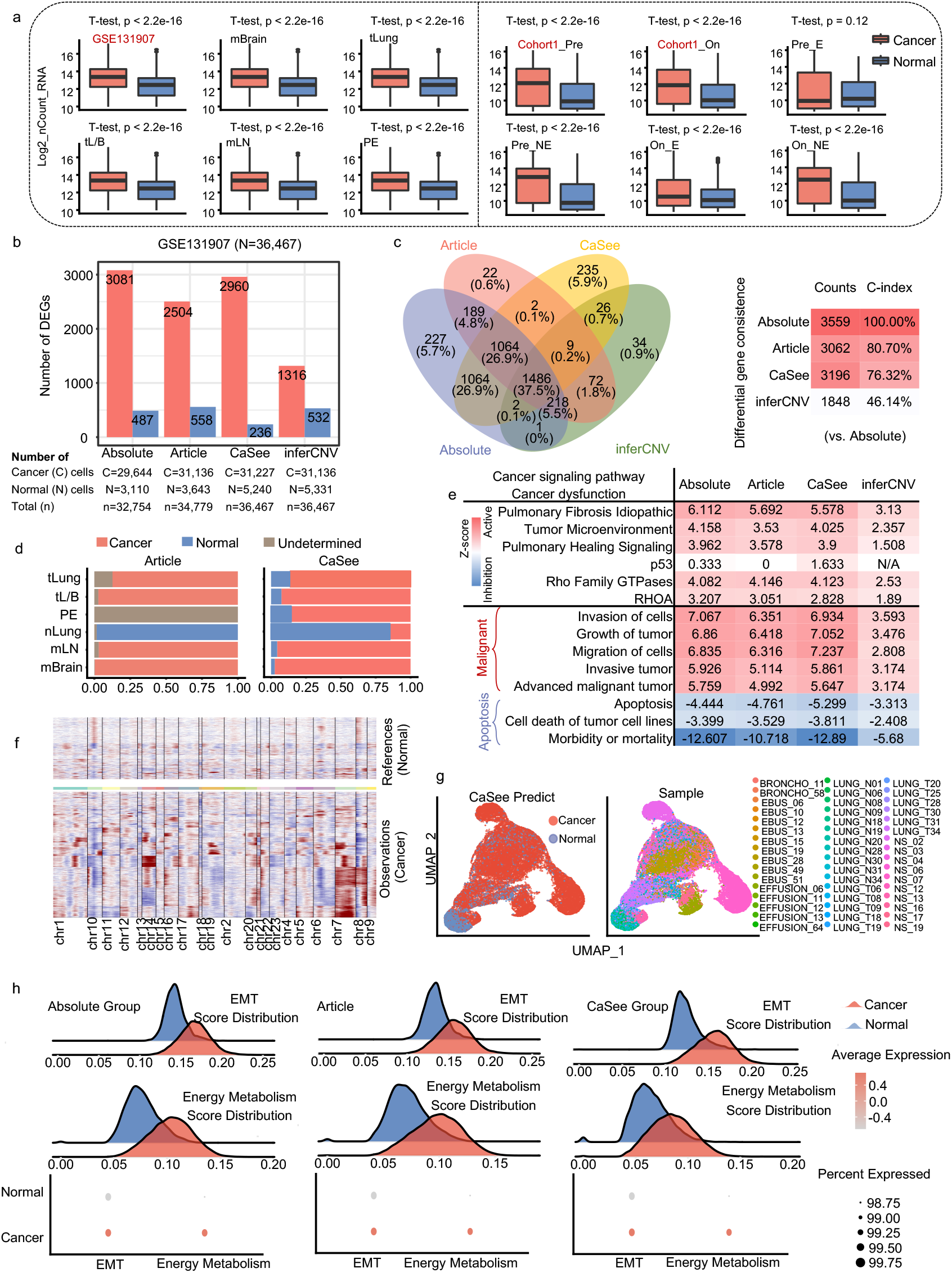
Comparison of differences between cancer cells and normal cells discriminated by different methods. Groups: Article, according to article provide cell subtype if cell state is nan or undetermined set as NA, in Article group without 1,688 NA cells; CaSee, comes from CaSee scale predict result; inferCNV, according to article provide cell subtype if cell state is nan or undetermined set as normal; Absolute, both Article and CaSee support the same cell state. A, Y-axis represent the number of gene counts (*log2*), X-axis represent CaSee discrimination result, red: Cancer, blue: Normal. B, The total number of cells in four groups and the number of DEGs in each group (cancer *vs*. normal). C, venn plot of four groups’ DEGs (right) and heatmap of the consistence of DEGs with absolute group (left). D, Cell distribution at different biopsy sites. E, IPA analysis of different method result, representative cancer signaling pathway (black line top) and cancer dysfunctional (black line bottom), z-score value represents the activation level of pathways/dysfunctional (z-score > 0, activation; z-score < 0, inhibition). F, CNV analysis of CaSee discriminated cells, the references are CaSee-normal cells, the observations are CaSee-cancer cells. G, left, UMAP of CaSee result. Right, UMAP of different samples. H, the module score distribution of Epithelial mesenchymal transformation (EMT) and energy metabolism related gene expression in cancer/normal cells discriminated by different methods.

Different from previous studies, we set homologous cells as references to infer the CNVs. In CaSee group, inferCNV analysis was carried out with normal cells as references and cancer cells as observations. It was found that some CNV noises existed in normal cells, but they are more in cancer cells, which are consistent with the previous hypothesis, that is, the genome of both cancer cells and normal cells in the same TME will change, and this change is more obvious in cancer cells. In other words, normal cells in the TME are “domesticated” by cancer cells, and the proportion of their own CNVs increases^13^ (Fig. 3F). Based on all the above, we need to note three main points. First, simply using inferCNV to discriminate cancer/normal cells will cause a controversial result. Second, when inferCNV considers normal cells (with elevated CNV levels) in the TME as references to identify cancer cells, errors will be introduced, leading to fewer cancer cells being identified (Fig. 2D PRJNA591860). Third, manually setting the CNV threshold could avoid the bad influence (Fig. 2D, GSE131907). However, CaSee could avoid this thorny problem because it is independent of those reference cells and directly identify the target cells.

Subsequently, we applied the Uniform Manifold Approximation and Projection (UMAP) method to reduce the dimension of the gene expression profile of candidate cancer cells. The results revealed that the heterogeneity between samples cannot be avoided by simply reducing the dimension of gene expression profiles. In other words, in the candidate cancer cell community, the heterogeneity between samples was much greater than that between cells (Fig. 3G). This conclusion is the same as Bischoff P.*et al*^37^ (Supplementary Fig. 2 and 3). Next, we calculated the score of EMT and energy metabolism modules (Supplementary material module_genes), respectively, using Ucell method^40^ to avoid the impact of data heterogeneity. The results showed that the activation levels of EMT and energy metabolism were significantly different in cancer and normal cells in Absolute group, Article group and CaSee group, and this trend was totally the same in the three groups (Fig. 3H). In general, whether in simple or complex research, CaSee performs well in the discrimination of candidate cancer cells.

### Application of CaSee in exploring the changes of cancer cells before and after treatment of PD-L1 breast cancer

We used the research data of Bassez A *et al*.^6^, to analyze cancer cell heterogeneity during PD-L1 treatment. Bassez A *et al*. used CD24, KRT19 and SCGB2A2 expression level, which are related to breast cancer to define cancer cells. We thought those cells just belong to candidate cancer cells, so CaSee appears. We evaluated the changes in the number of cancer/normal cells in expansion patients (E) and non-expansion patients (NE) before and on/after PD-L1 treatment. After receiving PD-L1 treatment, the proportion of cancer cells in type-NE patients was higher than that in type-E patients (*p*<0.05), which is the same as the conclusion drawn by Bassez A *et al*. (Fig. 4A, B). ANKRD30A and SYTL2 are marker genes of the luminal type. Combined with the cancer marker genes provided by Bassez A *et al*., we further evaluated DEGs between cancer and normal cells. The results showed that the five genes related to breast cancer had higher expression levels in cancer cells (Fig. 4C). According to the results of TCGA survival analysis, the high expression of these two genes affected the prognosis of patients (Fig. 4D). According to the expression level of gene markers, we further annotated the cell types of candidate cancer cells, including three main subtypes, namely, Basal_TP63 (+),Basal_TP63 (-) and luminal (Fig.4E). Basal_ TP63 (+) was mainly enriched in normal cells, whereas luminal cells were mainly enriched in cancer cells (Fig. 4F, *p*<0.05). The p63 is a myoepithelial cell marker and its low expression in cancer tissue is usually associated with the poor prognosis of breast cancer^41^. We performed IPA analysis on the two cohorts, respectively. However, due to the large difference in the number of cells between normal cells and cancer cells in cohort1, we did not obtain enough information on the status of pathways. It suggested that in this study, all samples were of high purity, but CaSee still found some activated cancer-related functions (Fig. 4G). In addition, we also performed IPA analysis on cohort2 (received neoadjuvant chemotherapy for 20–24 weeks)^6^ *vs*. cohort1 in all cancer cells. Interestingly, almost all cancer cell pathways and functions were significantly down-regulated in cohort2 before and on/after treatment, indicating that the treatment effect of combined PD-L1 after neoadjuvant chemotherapy is better (Fig. 4H).

**Fig. 4.**
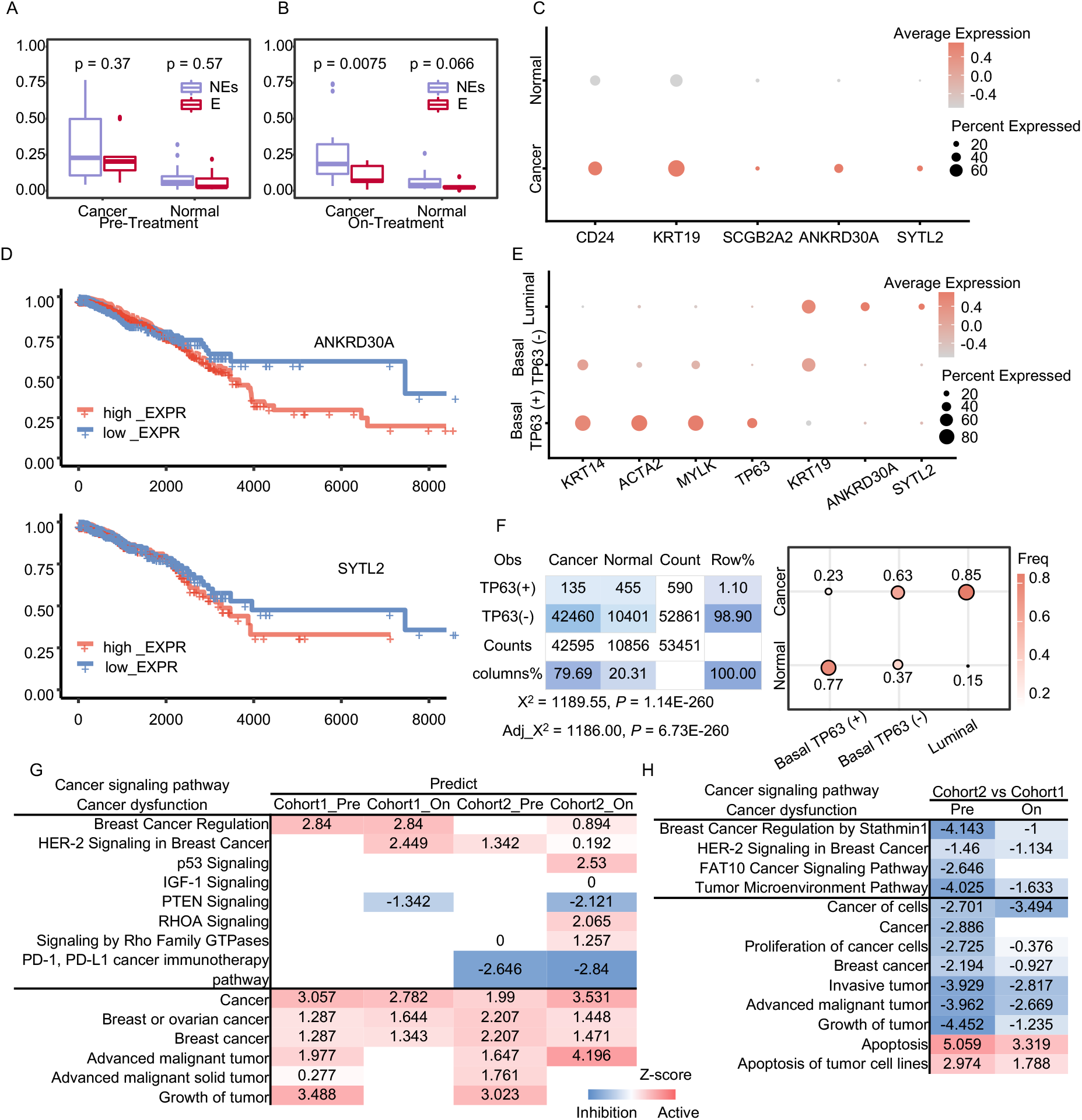
Practical application analysis of CaSee. A-B, Wilcoxon test applied in CaSee (pre-versus on-treatment). C, Average gene expression of selected marker genes for cancer and normal cells. D, Kaplan–Meier overall survival curves of GDC TCGA Breast Cancer (BRCA) (n = 1,284 samples)patients. +: censored observations. E, Average gene expression of selected marker genes for three types of candidate cancer cells. F, Fisher exact test P<0.0001, distribution of subtypes in cancer-normal cells. G-H, pathway and biofunction activation analysis using IPA in different cohort and time point (DEGs: cancer cell/normal cell)

## Discussion

Here, we established the CaSee pipeline to discriminate cancer/normal in the scRNA-seq expression matrix based on the transfer learning method. This pipeline is fast, accurate, efficient, and stable, and it is the first formal framework to directly solve the problem of cancer cell discrimination at the single-cell level. CaSee is a pipeline constructed based on the transfer learning method. It can recognize the feature space “knowledge” between cancer and normal cells in high-quality pan-cancer bulk-seq data, and successfully apply the “knowledge” into scRNA-seq data, to realize rapid and accurate pan-cancer cell discrimination. In cancer scRNA-seq, the identification of immune cells or stromal cells has relatively stable marker genes. However, it is difficult to quickly and accurately delineate malignant cancer cells to discriminate from scRNA-seq expression level owing to the different levels of heterogeneity caused by differences in samples, patients, and sequencing technologies. The feature extraction and routing mechanism in CaSee pipeline can extract the implicit feature space existing in the expression profile, which can help accurately and efficiently identify cancer cells. At the sequencing platform level, CaSee is robust and compatible with scRNA-seq technologies, such as 10x Genomics Chromium, Smart-seq2 or Microwell-seq^36^, and it will not be restricted by the data source. CaSee show strong discrimination ability in tissues, cell lines source or xenograft cells. However, traditional CNV detection methods depend on the data source. If there is no certain amount of reference cells in the data, the reliability of CNV detection methods will be greatly reduced. Different from the traditional CNV detection methods, CaSee is based on the deep learning method, so it can catch the “free ride” of GPU and realize ultrafast cell discrimination. In addition, CaSee pipeline shows its strong robustness in the evaluation of a variety of research data. In all studies, ultra-high similarity can be achieved. Compared with the traditional CNV detection methods that indirectly delineating cell types, direct discrimination of CaSee will bring researchers unparalleled data analysis experience. Nonetheless, CaSee might cause little errors, but these errors will not change the whole trend caused by the differences between cancer and normal cells, such as function/pathway module (Fig. 3E, H) and average gene expression (Supplementary Fig. 4).

In the data analysis of GSE131907, we found that the inferCNV method could indeed achieve accurate discrimination. However, it has strict requirements for data and after filtered by the threshold, some cells will be excluded. CaSee method can well address these two problems well. CaSee pipeline cannot replace the whole role of CNV methods that could calculate the genomic CNV in scRNA-seq expression matrix, but CaSee can completely and perfectly replace the role of cancer/normal cells discrimination, which could let each method performs its own functions.

A significant limitation of CaSee pipeline is that it is unable to accurately distinguish immune cells, fibroblast cells or other non-candidate cancer cells. Therefore, the type of input data for CaSee can only be candidate cancer cell expression matrix. Fortunately, except for candidate cancer cells, other cell types have relatively stable cell markers. Therefore, we have developed a “cell exclusion” method to obtain candidate cancer cells. We have already added this process into CaSee pipeline. Another limitation is that CaSee is mainly a black box developed based on capsule neural network, so it hardly to visualize the execution process. In addition, because the feature space of the “candidate reference data” is unchangeable (i.e., the number and type of genes are determined). considering that the intersection of the feature space in scRNA-seq data is too low to train. Therefore, we also open source the training tutorial of the whole model, so that users can select their own feature space and samples for training and transfer.

To sum up, CaSee provides a powerful automatic method for faster, better, and easier cancer/normal cells discrimination. It is suitable for most sequencing technologies and data sources and is widely used in various cancer origins. CaSee has completely changed the link between bulk-seq and scRNA-seq and built a new connecting bridge. It can more accurately identify the differences between cancer and normal cells at the scRNA-seq level, which helps us further understand the heterogeneity and homogeneity between various cancer cells. With the rapid development of deep learning, the field of life science has entered a new era of digital intelligence. Making good use of CaSee will greatly improve our research methods and our cognitive level of cancer.

## Methods

### TCGA and GTEx Data process

We’re from UCSC Xena (https://xenabrowser.net/datapages/) Download TCGA and GTEX data, using R package org.hs.eg.db (10.18129/B9.bioc.org.Hs.eg.db) to switch gene ID to gene symbol. The number of rare cancer types may affect the training process of the model. Therefore, the number of cancer type in the candidate reference dataset should be greater than 0.5% total samples (supplementary Table 1). In all control samples, TCGA samples accounted for 10% of the total. Code: https://github.com/yuansh3354/CaSee/Get_Normal_sample_candidate_cancer_cells.r

### TCGA survival analysis

Survival analysis was executed in UCSC Xena, using GDC TCGA Breast Cancer (BRCA) HTSeq-FPKM gene expression profile. ANKRD30A gene expression cut-off is 0.6; SYTL2 using quartiles cut-off. Survival curves were fitted using the Kaplan–Meier formula in the R package ‘survival’ and visualized using the ggsurvplot function of the R package ‘survminer’. Code: https://github.com/yuansh3354/CaSee/blob/main/Survival%20analysis.R

### CTC isolation and single cell RNA sequence

Pleural effusion sample was collected from a breast cancer patient from the Fifth Medical Center of Chinese People’s Liberation Army General Hospital. First, the pleural effusion sample was centrifuged at 1500 rpm to remove the supernatant. Then, 5mL of red blood cell lysing buffer (Beyotime) was added to lyse red blood cells for 15 min. The nucleated cells were pelleted and resuspended in PBS. The cells were stained with FITC-conjugated anti-EpCAM (abcam), CY5-conjugated anti-HER2 (Biolegend) and PE-conjugated anti-CD45 (abcam). Single tumor cells (EpCAM-Positive/HER2-Positive/CD45-Negative) were sorted out by FACS. Finally, the single tumor cells were treated following the protocol of SMART-Seq v4 Ultra Low Input RNA Kit (Takara) for single-cell RNA sequence. And ERCC was added to each sample as a known amount of exogenous intrusion.

### Capsule Network Training loop

Feature “extraction-integration” network structure: the input scRNA-seq 1D tensor data is passed through the layer convert into 2D feature map tensor (size 28*28). Each pixel unit in the feature map can be directly related to a gene pattern (nonlinear combination of multiple genes). The structure consists of a fully connected layer with bias and a ReLU activation function. Primary capsule layer: the network mainly performs convolution functions. This network layer is composed of 32 main capsules, which compresses and outputs the gene module characteristics in the convolution layer and transmits them to the digital capsule layer. Digital capsule layer: this capsule layer is mainly used for capsule integral calculation, and finally outputs the probability of “normal_like” and “tumor_like” respectively (independently). Capsule reconstruction layer: decode and reconstruct the output of digital capsule layer to recover the original data as much as possible. Its main function is to maintain the feature space.

Training loop

Step_0_Data_Prepare

1. Users can input raw count matrix and give the marker genes of T cell (default CD3D, CD3E, CD2), Fibroblast cell (default COL1A1, DCN, C1R), myeloid cell (default LYZ, CD68, TYROPB), B cell (default CD79A, MZB1, MS4A1), endothelia cell (default CLDN5, FLT1, RAMP2), mast cell (default CPA3, TPSAB1, TPSB2), DC cell (default LILRA4, CXCR3, IRF7), and candidate cancer cell (default EPCAM or NA). If users input raw count matrix, CaSee pipeline will run the stander scRNA-seq data process protocols in Scanpy to do cell clusters annotation. Then remain the candidate cancer cell matrix to do training loop.
2. Users can input candidate cancer cells count matrix. Then all cell annotation pipelines will be skipped.

Step_1_Training_Loop: First, the candidate cancer cell count matrix will intersect with the candidate reference data to generate reference data; Second, the reference data split into training data, validation data and test data (8:1:1). The training data mainly participate in the parameter training of the model, the verification data is responsible for controlling the model, and the test data is used to evaluate the classification performance of the final model. When the accuracy of the test data can reach more than 95%, training loops finished. Finally, use the model into scRNA-seq candidate cancer cell expression matrix.

### Model Transfer and Discriminating Single-Cell Data

Capsule neural network is a kind of multi-label and multi-classification model, so the output result is NormalLike and TumorLike independent of each other. Therefore, we corrected the output results in the process of scRNA-seq discrimination. The formulas as:

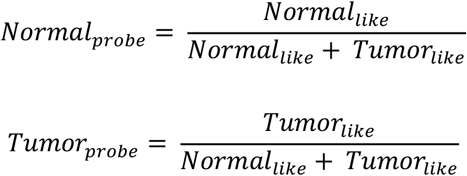

In addition, considering the heterogeneity between different samples and the purity of cancer cells, we also designed a weighted correction probability. The weighted correction probability is usually applied to data from the same point in time or the same patient. Typically, for example, CTC scRNA-seq data, cell line scRNA-seq data, and multiple sampling scRNA-seq data at different locations of a single patient. The weighting calculation formulas as:

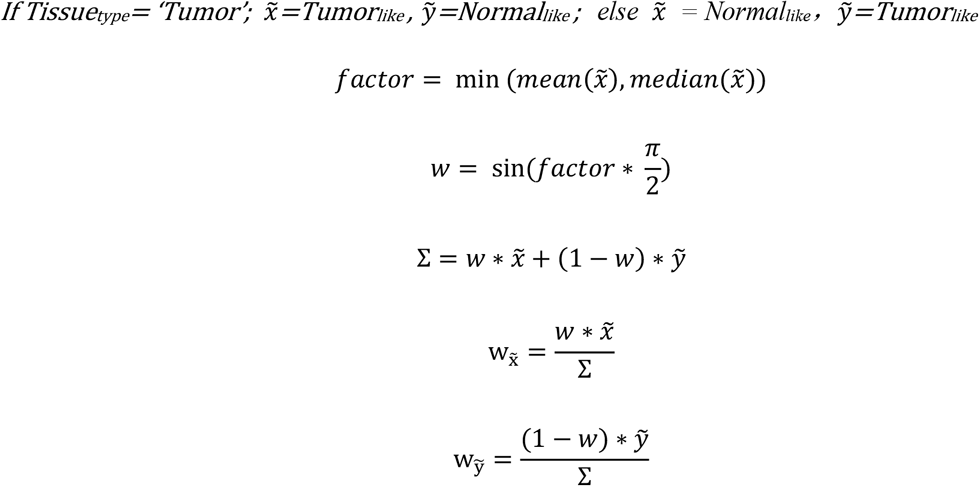

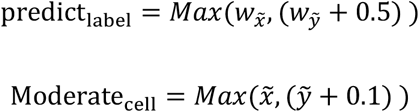

Users can use the multiple-cross verification (10 times by default, usually 1∼1.5 minutes / per cross) to ensure the stability of discriminate results. All the model output results in this research were verified by 10x-cross verification, the p-value of each cell was calculated by cumulative binomial distribution test, and the p-value was corrected by BH method. p-value calculated as follow:

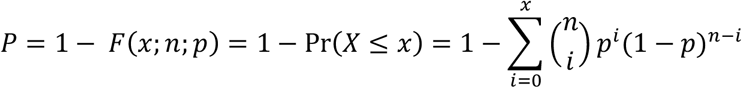

Generally, the probability of cancer cells and normal cells are equal, P = 0.5

### scRNA-seq data processing

We considered all cell line sources and xenograft cell sources as cancer cells. Research based on tissue sources we considered epithelial cells as candidate cancer cells. We considered Parietal, Gastric, Pit, Epithelial, AT2, AT1, Clara, and Hepatocyte cell as candidate cancer cell in normal tissue source (see Code:https://github.com/yuansh3354/CaSee/blob/main/Get_Normal_sample_candidate_cancer_cells.r). The “cell exclusion” algorithm had been integrated in the CaSee pipeline. If the user chooses to input the raw count expression matrix, the algorithm will start automatically. Users can set marker gene lists of different cell types according to their own research background (as above). The cell exclusion algorithm is based on Python 3.8 8,Scanpy 1.7. 2. Firstly, the algorithm filters high variable genes (top 2000) from the count expression matrix entered by the user, and then executes scale (default parameter), PCA (default parameter), neighbors (default parameter) and dimension (default parameter) in turn. The first 20 PCA were used for tSNE. According to the marker gene list inputted by the user, different cell clusters are labeled respectively, and the candidate cancer cell gene expression matrix is output.

### Hardware and software information

The configuration of the personal computer accessories is as follows (There are no overclocking Settings):CPU, AMD3950X; GPU, NVIDIA GeForce 3090; Memory, DDR4-3200 32*4; operating system environment, Ubuntu 20.04 LTS. CaSee is developed based on Python version 3.8.8 The main modules used include Python version 1.9.0, cuda11 and python lightning version 1.3.7. R version 4.1.0. The main modules used include Seurat version 4.0.4, CopyKAT version 1.0.4 and inferCNV version 1.8.1. See the configuration table for detailed parameters.

### Ingenuity Pathway Analysis

Ingenuity Pathway Analysis (IPA) is a bioinformatics analysis method. We use IPA method to locate features and annotate functions. P-value <0.05 was considered a statistically significant threshold. Z-value greater than 0 is defined as active, and less than 0 is defined as suppressed. The activation z-score of a hypothesis is calculated from the regulation directions and gene expression changes of the genes in the overlap of data set and hypothesis-regulated genes. It assesses whether there is a significant pattern match between predicted and observed up- and down-regulation, and also predicts the activation state of the regulator (z > 0: activating, z < 0: inhibiting). The activation z-score is given by

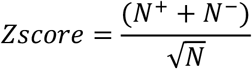

with *N*^*+*^(*N*^*-*^) being the number of genes where the product of net-effect and observed direction of gene regulation is greater (less) than zero, and *N = N*^*+*^ *+ N*^-42^.

## Data and materials availability

All data download address links used in this study are in supplementary Table 2. Raw CTC single-cell sequenceing data is available in the China National Center for Bioinformation (BioProject ID PRJCA007531). All data processing codes, and configuration files are stored on GitHub (https://github.com/yuansh3354/CaSee).

## Ethics approval and Consent to participate

Not applicable

## Consent for publication

Not applicable

## Contributions

X.Z, Y.S. and Z.H. designed the study. Y.S. and C.G. performed the analyses and interpreted the results. Y.S and X.Z wrote the manuscript. F.S and F.J. performed CTC isolation. X.Z. conducted this study. All authors read and approved the final manuscript.

## Corresponding authors

Correspondence to Xiuli Zhang, Zhiyuan Hu.

## References

1. Wagner, J. et al. A Single-Cell Atlas of the Tumor and Immune Ecosystem of Human Breast Cancer. Cell 177, 1330-1345.e18 (2019).

2. Villani, A.-C. et al. Single-cell RNA-seq reveals new types of human blood dendritic cells, monocytes, and progenitors. Science 356, eaah4573 (2017).

3. Peng, J. et al. Single-cell RNA-seq highlights intra-tumoral heterogeneity and malignant progression in pancreatic ductal adenocarcinoma. Cell Res 29, 725–738 (2019).

4. Patel, A. P. et al. Single-cell RNA-seq highlights intratumoral heterogeneity in primary glioblastoma. Science 344, 1396–1401 (2014).

5. Nguyen, Q. H. et al. Profiling human breast epithelial cells using single cell RNA sequencing identifies cell diversity. Nat Commun 9, 2028 (2018).

6. Bassez, A. et al. A single-cell map of intratumoral changes during anti-PD1 treatment of patients with breast cancer. Nat Med 27, 820–832 (2021).

7. Kim, N. et al. Single-cell RNA sequencing demonstrates the molecular and cellular reprogramming of metastatic lung adenocarcinoma. Nat Commun 11, 2285 (2020).

8. Zhang, X. et al. CellMarker: a manually curated resource of cell markers in human and mouse. Nucleic Acids Research 47, D721–D728 (2019).

9. Yuan, H. et al. CancerSEA: a cancer single-cell state atlas. Nucleic Acids Research 47, D900–D908 (2019).

10. Oh, D. Y. et al. Intratumoral CD4+ T Cells Mediate Anti-tumor Cytotoxicity in Human Bladder Cancer. Cell 181, 1612-1625.e13 (2020).

11. Taylor, A. M. et al. Genomic and Functional Approaches to Understanding Cancer Aneuploidy. Cancer Cell 33, 676-689.e3 (2018).

12. Zarrei, M., MacDonald, J. R., Merico, D. & Scherer, S. W. A copy number variation map of the human genome. Nat Rev Genet 16, 172–183 (2015).

13. Zhou, Y. et al. Single-Cell Multiomics Sequencing Reveals Prevalent Genomic Alterations in Tumor Stromal Cells of Human Colorectal Cancer. Cancer Cell 38, 818-828.e5 (2020).

14. Gao, R. et al. Delineating copy number and clonal substructure in human tumors from single-cell transcriptomes. Nat Biotechnol 39, 599–608 (2021).

15. Shao, X. et al. scDeepSort: a pre-trained cell-type annotation method for single-cell transcriptomics using deep learning with a weighted graph neural network. Nucleic Acids Research 49, e122–e122 (2021).

16. He, Y., Yuan, H., Wu, C. & Xie, Z. DISC: a highly scalable and accurate inference of gene expression and structure for single-cell transcriptomes using semi-supervised deep learning. Genome Biol 21, 170 (2020).

17. Yamada, H. et al. Predicting Materials Properties with Little Data Using Shotgun Transfer Learning. ACS Cent. Sci. 5, 1717–1730 (2019).

18. Zhu, R. et al. Phase-to-pattern inverse design paradigm for fast realization of functional metasurfaces via transfer learning. Nat Commun 12, 2974 (2021).

19. Hu, J. et al. Iterative transfer learning with neural network for clustering and cell type classification in single-cell RNA-seq analysis. Nat Mach Intell 2, 607–618 (2020).

20. Qiu, Y. L., Zheng, H., Devos, A., Selby, H. & Gevaert, O. A meta-learning approach for genomic survival analysis. Nat Commun 11, 6350 (2020).

21. Bell, C. C. et al. Targeting enhancer switching overcomes non-genetic drug resistance in acute myeloid leukaemia. Nat Commun 10, 2723 (2019).

22. Maynard, A. et al. Therapy-Induced Evolution of Human Lung Cancer Revealed by Single-Cell RNA Sequencing. Cell 182, 1232-1251.e22 (2020).

23. Roels, J. et al. Distinct and temporary-restricted epigenetic mechanisms regulate human αβ and γδ T cell development. Nat Immunol 21, 1280–1292 (2020).

24. Xi, E., Bing, S. & Jin, Y. Capsule Network Performance on Complex Data. 1712.03480 [cs, stat] (2017).

25. Wang, L. et al. An interpretable deep-learning architecture of capsule networks for identifying cell-type gene expression programs from single-cell RNA-sequencing data. Nature Machine Intelligence 2, 693–703 (2020).

26. Qiao, K. et al. Accurate reconstruction of image stimuli from human fMRI based on the decoding model with capsule network architecture. 14.

27. Krizhevsky, A., Sutskever, I. & Hinton, G. E. ImageNet classification with deep convolutional neural networks. Commun. ACM 60, 84–90 (2017).

28. Sabour, S., Frosst, N. & Hinton, G. E. Dynamic Routing Between Capsules. 1710.09829 [cs] (2017).

29. Kharchenko, P. V., Silberstein, L. & Scadden, D. T. Bayesian approach to single-cell differential expression analysis. Nat Methods 11, 740–742 (2014).

30. Rambow, F. et al. Toward Minimal Residual Disease-Directed Therapy in Melanoma. Cell 174, 843-855.e19 (2018).

31. Kim, N. et al. Single-cell RNA sequencing demonstrates the molecular and cellular reprogramming of metastatic lung adenocarcinoma. Nat Commun 11, 2285 (2020).

32. Lee, H.-O. et al. Lineage-dependent gene expression programs influence the immune landscape of colorectal cancer. Nat Genet 52, 594–603 (2020).

33. Gao, R. et al. Delineating copy number and clonal substructure in human tumors from single-cell transcriptomes. Nat Biotechnol (2021) doi:10.1038/s41587-020-00795-2.

34. Oren, Y. et al. Cycling cancer persister cells arise from lineages with distinct programs. Nature 596, 576–582 (2021).

35. Maynard, A. et al. Therapy-Induced Evolution of Human Lung Cancer Revealed by Single-Cell RNA Sequencing. Cell 182, 1232-1251.e22 (2020).

36. Han, X. et al. Construction of a human cell landscape at single-cell level. Nature 581, 303–309 (2020).

37. Bischoff, P. et al. Single-cell RNA sequencing reveals distinct tumor microenvironmental patterns in lung adenocarcinoma. Oncogene (2021) doi:10.1038/s41388-021-02054-3.

38. Maynard, A. et al. Therapy-Induced Evolution of Human Lung Cancer Revealed by Single-Cell RNA Sequencing. Cell 182, 1232-1251.e22 (2020).

39. Zhang, L. et al. Gene Expression Profiles in Normal and Cancer Cells. Science 276, 1268–1272 (1997).

40. Andreatta, M. & Carmona, S. J. UCell: Robust and scalable single-cell gene signature scoring. Computational and Structural Biotechnology Journal 19, 3796–3798 (2021).

41. Hu, L. et al. ΔNp63α is a common inhibitory target in oncogenic PI3K/Ras/Her2-induced cell motility and tumor metastasis. Proc Natl Acad Sci USA 114, E3964–E3973 (2017).

42. Krämer, A., Green, J., Pollard, J. & Tugendreich, S. Causal analysis approaches in Ingenuity Pathway Analysis. Bioinformatics 30, 523–530 (2014).

